# Spectraplakin is required for dendritic microtubule organization via a tip-based mechanism

**DOI:** 10.1101/2025.06.27.661943

**Authors:** Matthew Davies, Neeraja Sanal, Neele Wolterhoff, Ulrike Gigengack, Yitao Shen, Samuel Matthew Frommeyer, Ines Hahn, Sebastian Rumpf

## Abstract

The differential microtubule organization in axons and dendrites underlies neuronal polarity and developmental processes like neurite pruning. How neurons achieve their specific microtubule organization during development is an area of active research, and it has been especially hard to explain how dendritic microtubules are oriented with their plus ends towards the soma. Transient microtubule nucleation from tips of early growing dendrites has been detected in some systems and would explain how orientation is set up. In a survey for cytoskeletal regulators involved in dendrite pruning and microtubule organization in Drosophila, we found the spectraplakin Short stop (Shot), an actin/microtubule crosslinker. Loss of Shot causes microtubule orientation defects already during early dendrite development, when Shot is transiently recruited to tips of growing dendrites via its actin binding domain. Genetic and functional evidence suggest that Shot’s primary function in this process is to locally stabilize microtubules. We also provide evidence for a developmentally transient microtubule nucleation mechanism. Our data highlight the importance of transient and localized microtubule regulation for dendritic microtubule organization.

## Introduction

Microtubules are long polar tube-like polymers composed of α/β-tubulin dimers that provide mechanical stability to cells and act as tracks for motor-based transport. Tubulin dimers are preferentially added, and also lost, at microtubule plus ends, whereas microtubule minus ends are relatively inert and often localize to microtubule organizing centers (MTOCs), structures of diverse composition that nucleate or anchor microtubules. In dividing cells, the main MTOCs are centrosomes, but many non-dividing or differentiated cells have less well characterized non-centrosomal (nc)MTOCs (Akhmanova & Kapitein, 2022).

Given their intrinsic polarity, microtubules, and the way they are organized in a given cell, play an important role in cellular polarity. This is particularly evident in neurons. In axons, all microtubules are oriented with their plus ends towards the distal synapses, thus enabling axon-specific transport, especially via kinesin-1. In contrast, a large fraction of dendritic microtubules is oriented with the plus ends towards the soma (“plus end-in”), enabling dendrite-specific transport via other motors including dynein. The plus end-in orientation is especially prevalent in invertebrates, where microtubules in primary dendrite branches are often almost uniformly plus end-in (Rolls *et al*, 2007). Despite the recognized importance of microtubule organization for neuronal function, it is still poorly understood how neuronal - and in particular, dendritic - microtubule organization is set up and maintained.

The differential organization of neuronal microtubules also underlies important neuronal morphogenetic processes. For example, the peripheral nociceptive class IV dendritic arborization (c4da) neurons of *Drosophila* larvae prune their long and branched sensory dendrites at the onset of metamorphosis. Pruning involves severing of proximal dendrite regions and subsequent degeneration of distal fragments (Rumpf *et al*, 2017; Furusawa & Emoto, 2021). The presumptive severing sites are marked by the local loss of microtubules, which predisposes them to mechanical tearing (Krämer *et al*, 2023). This local microtubule disassembly is initiated by an increase in microtubule dynamics at the onset of the pupal stage (Herzmann *et al*, 2017). Due to their uniform plus end-in orientation, the dendritic microtubules shrink from their plus ends in a coordinated fashion, eventually leading to the complete loss of microtubules in proximal dendrite regions (Herzmann *et al*, 2018; Rumpf *et al*, 2019). In support of this model, conditions that cause mixed orientation of dendritic microtubules either cause pruning defects or act as genetic enhancers of pruning mutants with microtubule-related functions. For example, the motor protein kinesin-2 together with its binding partner, the plus end-binder End-Binding protein 1 (EB1), guide polymerizing microtubules along preexisting ones, thus ensuring the maintenance of plus end-in orientation (Mattie *et al*, 2010). Loss of kinesin-2 causes pruning defects, while EB1 knockdown genetically enhances the pruning defects caused by the tau kinase Par-1 (Herzmann *et al*, 2018). Dendritic microtubule orientation and pruning defects are also caused by loss of the minus end binding protein Patronin (Wang *et al*, 2019) and the serine/threonine phosphatase PP2A (Rui *et al*, 2020; Wolterhoff *et al*, 2020). Loss of the microtubule polymerase Minispindles (Msps)/XMAP-215 (Tang *et al*, 2020) also causes defects in microtubule orientation, possibly linking dendritic microtubule orientation to microtubule growth or nucleation.

While several factors are known to maintain uniform dendritic microtubule orientation in Drosophila peripheral neurons, it is not clear how dendritic microtubule orientation arises during development. One model for this has been put forward based on the *C*. *elegans* PVD neuron model system. Here, a ncMTOC residing on Rab11-positive recycling endosomes is localized at the tip of the outgrowing dendrite, such that microtubules can only grow towards the soma, i. e., plus end-in (Liang *et al*, 2020; Harterink *et al*, 2018). Whether tip-based mechanisms could also inform dendritic microtubule polarity in other systems is not known.

Spectraplakins are a family of actin/microtubule cytoskeletal crosslinker proteins with important roles in microtubule organization in differentiated cells (Sanchez *et al*, 2021; Sun *et al*, 2019; Voelzmann *et al*, 2017). Anchored to the cell cortex or other actin-rich structures via an N-terminal actin binding domain, they can tether microtubules via binding sites for the minus end binding protein Patronin (Noordstra *et al*, 2016; Ning *et al*, 2016; Nashchekin *et al*, 2016), furthermore, spectraplakins can recruit factors related to microtubule growth such as the plus end binder EB1 (Slep *et al*, 2005). A neuronal function has been shown for the *Drosophila* spectraplakin Short stop (Shot), which bundles and stabilizes microtubules in axons (Alves-Silva *et al*, 2012, Okenve-Ramos *et al*, 2024). This effect is mediated by its C-terminal GAS2 domain (Okenve-Ramos *et al*, 2024), a type of microtubule-binding domains known to bundle and sometimes even nucleate microtubules (An *et al*, 2025).

Here, we describe an important role for Shot in dendritic microtubule organization. Loss of Shot in c4da neurons causes mixed orientation of dendritic microtubules and consequently, dendrite pruning defects. We find that Shot is already required for uniform orientation during early dendrite growth stages, when it transiently localizes to dendrite tips in an actin-dependent manner. Domain analyses show that this actin-dependent localization is required for Shot’s function. The GAS2 domain is also required, and Shot can increase dendritic microtubule density, indicating that it contributes to dendritic microtubule organization via a local stabilization mechanism. We also provide evidence that Rab11 is important for dendritic microtubule orientation and might be part of a transient dendritic MTOC. Our data suggest that tip-based mechanisms could be broadly involved in dendritic microtubule organization and identify Shot as an important player.

## Results

### Shot is required for c4da neuron dendrite pruning

The long and branched dendrites of third instar larval c4da neurons are completely pruned at the onset of the pupal stage, such that by 16 hours after puparium formation (h APF), they have lost all their dendrites (Fig. 1 A, 1 B, B’). In order to identify new regulators of c4da neuron dendrite pruning, we expressed candidate RNAis in c4da neurons under *pickpocket (ppk)-GAL4* and assessed their effects at 16 h APF. This approach identified a dsRNA line targeting *Shot*, which encodes the sole Drosophila member of the spectraplakins, a family of large adaptor proteins with binding sites for actin and microtubules. C4da neurons expressing *shot* dsRNA had long and branched dendrites at the larval stage (Fig. 1 C), even though their dendritic field coverage seemed somewhat reduced compared to control neurons. At 16 h APF, dendrites could still be seen attached to the soma in almost half the c4da neurons expressing *shot* dsRNA (Fig. 1 C’, H - J), indicative of pruning defects. In order to confirm this result, we used Mosaic Analysis with a Repressible Cell Marker (MARCM) to generate c4da neuron clones homozygous for the loss-of-function allele *shot^3^* (Kolodziej *et al*, 1995). *shot^3^* mutant c4da neurons had shorter dendrites at the L3 stage compared to control MARCM c4da neuron clones (Fig. 1 D, E). At 16 h APF, most *shot^3^* mutant neurons still had dendrites attached to the soma (Fig. 1 D’, E’, H - J). We also generated an sgRNA line targeting an exon common to all Shot splice isoforms and found that it caused significant c4da neuron dendrite pruning defects when expressed in c4da neurons in a sensitized *shot^3^*/+ heterozygous background (Fig. 1 F’, G’, H - J).

**Figure 1.**
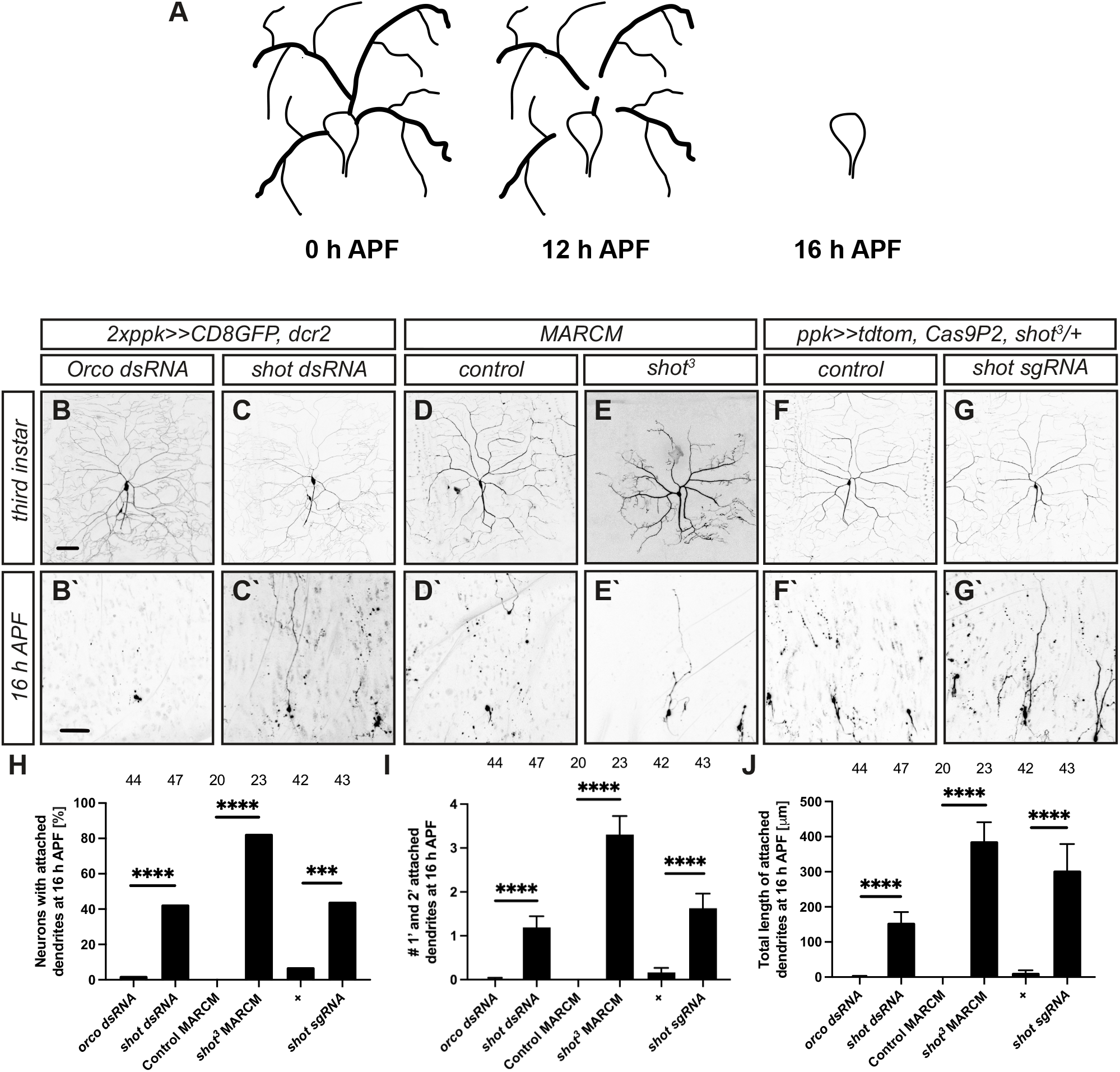
Shot is required for c4da neuron developmental dendrite pruning. **A** Schematic showing the progression of c4da neuron dendrite pruning via hormonal induction (0 h APF) to severing (12 h APF) and clearance (16 h APF). **B** – **G’** C4da neurons of the indicated genotypes were imaged at the third instar larval stage (**B** - **G**) and at 16 hours after puparium formation (h APF) (**B’ - G’**). **B, B’** Control c4da neurons expressing Orco dsRNA under the control of *ppk-GAL4*. **C, C’** C4da neurons expressing Shot dsRNA under the control of *ppk-GAL4*. **D, D’** Control c4da neurons labeled by MARCM. **E, E’** *shot^3^*mutant c4da neurons labeled by MARCM. **F, F’** Control c4da neurons expressing Cas9P2 under *ppk-GAL4* in a *shot^3^* heterozygous background. **G, G’** C4da neurons coexpressing Cas9P2 and Shot sgRNA under the control of *ppk-GAL4* in a *shot^3^* heterozygous background. **H** Phenotypic penetrance of pruning defects in **B’ - G’**. Number of neurons for each sample are indicated above the graph. *** P<0.001, **** P<0.0001, two-tailed Fisher’s exact test. **I** Average number of primary and secondary dendrites attached to the soma at 16 h APF in samples **B’ - G’**. Values are mean +/- s. e. m., **** P<0.0001, Mann-Whitney U test. **J** Total length of remaining dendrites at 16 h APF in samples **B’ - G’**. Values are mean +/- s. e. m., **** P<0.0001, Mann-Whitney U test. Scale bars in **B** and **B’** are 50 µm.

C4da neurons undergo extensive remodeling during the pupal stage. Concomitantly with their dendrites, they also prune their presynapses in an ecdysone- and actin-dependent manner (Furusawa *et al*, 2023; Frommeyer *et al*, 2026). Later during the pupal stage, c4da neurons grow new dendrites adapted to the adult stage (Sanal *et al*, 2023). While Shot was not required for presynapse pruning (Figure S1 A), c4da neurons lacking Shot also failed to initiate dendrite regrowth at 72 h APF, when control c4da neurons had regrown long and branched dendrites (Figure S1 B, C). We conclude that Shot is required for c4da neuron dendrite remodeling, but does not seem to affect synapses.

### Shot is required for uniform dendritic microtubule orientation

We and others have previously shown that c4da neuron dendrite pruning depends on local microtubule disassembly in proximal dendrites (Williams *et al*, 2005, Herzmann *et al*, 2017, Herzmann *et al*, 2018, Wang *et al*, 2019). To assess a potential effect of Shot knockdown on dendritic microtubule regulation during the early pupal stage, we visualized dendritic microtubules at 6 h APF using immunofluorescence against the microtubule-associated protein futsch/MAP1B. While control neurons consistently showed gaps in their futsch signal in these regions (Fig. 2 A, C), Shot knockdown neurons often still displayed continuous futsch staining in their proximal dendrites (Fig. 2 B, C).

**Figure 2.**
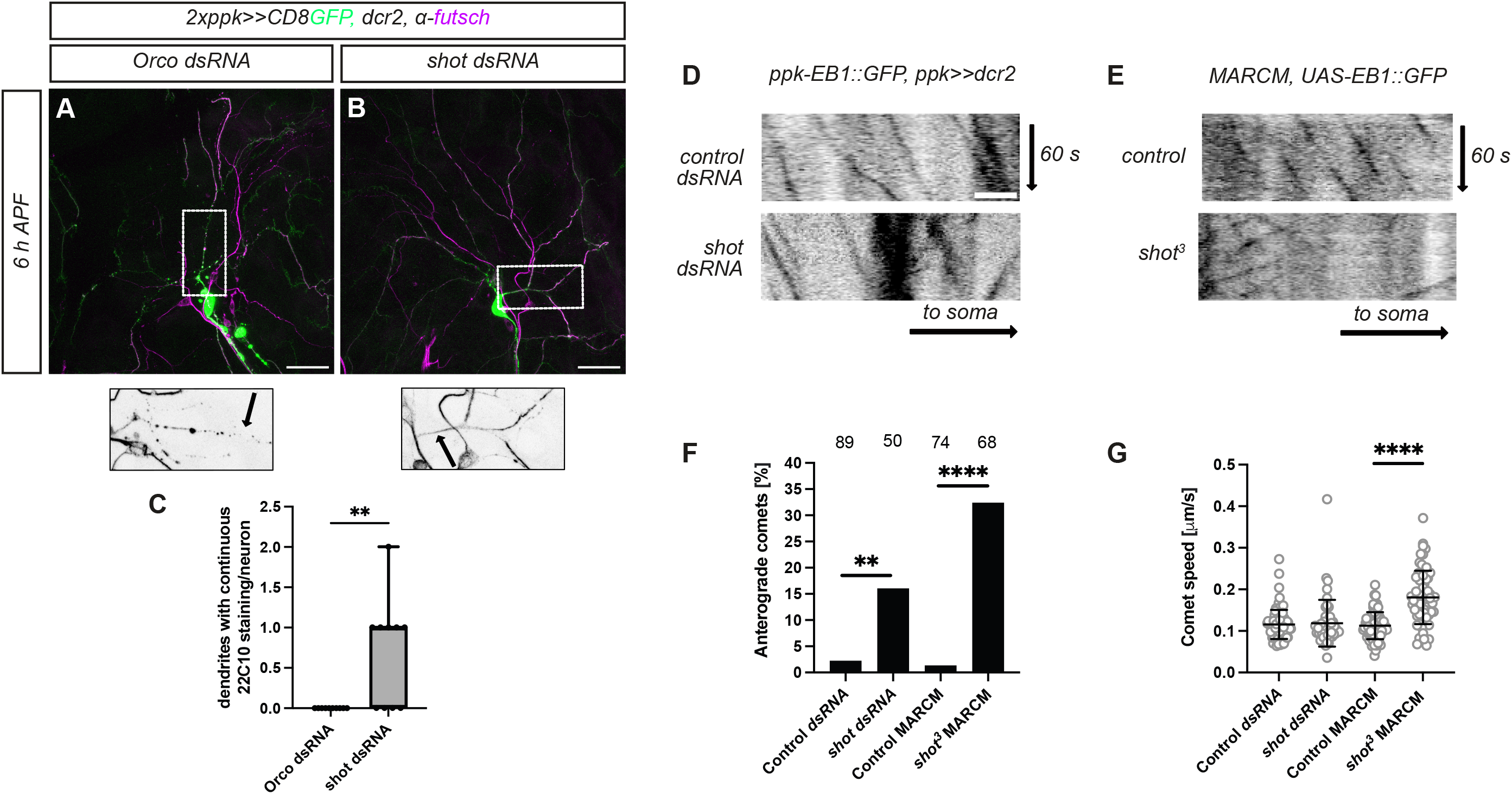
Shot is required for uniform plus end-in orientation of dendritic microtubules. **A** - **C** Delayed microtubule disassembly in neurons lacking Shot. Microtubules in control or Shot knockdown c4da neurons (green) were visualized by futsch/22C10 staining (magenta) at 6 h APF. Smaller panels below show futsch/22C10 signal in boxed areas in **A**, **B**, arrows indicate c4da neuron dendrites. **A** Control c4da neuron. **B** C4da neuron expressing Shot dsRNA. **C** Graph depicting the average number of dendrites with continuous futsch/22C10 staining per neuron in **A**, **B**. Box represents s. d., N = 10 neurons each, ** P<0.01, student’s t-test. **D**, **E** EB1::GFP comets were imaged in primary dendrites of third instar larval c4da neurons, and comet movement was depicted in kymographs. Time and direction of the soma are indicated by arrows. **D** Upper panel, EB1::GFP comets in control neuron expressing Orco dsRNA; lower panel, EB1::GFP comets in neuron expressing Shot dsRNA. **E** Upper panel, EB1::GFP comets in control c4da neuron MARCM clone; lower panel, EB1::GFP comets in *shot^3^*mutant c4da neuron MARCM clone. **F** Percentage of anterogradely moving comets in panels **D**, **E**. Numbers of comets per sample are indicated above the graph. ** P<0.01, **** P<0.0001, two-tailed Fisher’s exact test. **G** EB1::GFP comet speed in panels **D**, **E**. **** P<0.0001, Wilcoxon’s test. Scale bars in **A** and **D** are 20 and 5 µm, respectively.

As microtubules shrink away from the cell body in a coordinated fashion during dendrite pruning, the uniform plus end-in orientation of dendritic microtubules is particularly important for dendritic microtubule disassembly (Herzmann *et al*, 2018; Rumpf *et al*, 2019). To assess whether loss of Shot affects dendritic microtubule orientation, we expressed EB1::GFP in c4da neurons and live imaged it in dendrites. EB1 binds to the plus ends of growing microtubules (both newly nucleated and re-elongating), and plus end-bound EB1 is visible as moving dots (also known as comets). The movement direction of EB1 comets then indicates the orientation of the respective microtubule. As expected, EB1::GFP comets in primary or secondary dendrites of third instar control c4da neurons moved almost exclusively retrogradely towards the soma, indicating plus end-in orientation (Fig. 2 D, F). Upon *shot* knockdown, a significantly higher proportion of EB1::GFP comets could be seen moving anterogradely, or plus end-out (Fig. 2 D, F). To confirm this result, we assessed EB1::GFP behavior in *shot^3^* c4da neuron MARCM clones and observed an even higher proportion of plus end-out dendritic microtubules in comparison to control c4da neurons MARCM clones (Fig. 2 E, F). Comets in *shot^3^* mutant neurons also moved at a slightly higher speed than in control neurons (Fig. 2 G). We conclude that Shot is required for the correct orientation of dendritic microtubules.

### Actin and microtubule binding by Shot are crucial for pruning and microtubule organization

In order to better understand the function of Shot in dendritic pruning and microtubule orientation, we next asked which domains of Shot are crucial for these functions. Of particular interest were the domains of Shot that bind to actin and microtubules. Shot has two Calponin Homology (CH) domains at the N-terminus that mediate binding to specific actin structures, and a microtubule-binding GAS2 domain in its C-terminal region (Fig. 3 A). We next expressed full length or truncated Shot variants in *shot^3^* mutant c4da neurons and asked whether they could rescue the pruning and microtubule orientation defects. We used a genomic Shot BAC rescue construct with a C-terminal YFP-tag and a UAS construct containing GFP-tagged full length Shot UAS (Shot^FL^::GFP) as full length Shot variants. To address the role of the actin binding domain, we tested Shot^ΔCH1^::GFP, a Shot variant lacking the first CH domain with decreased affinity for actin (Lee & Kolodziej, 2002), and Shot^ΔABD^::GFP, which lacks both CH domains. To address the function of microtubule binding, we either used a full length Shot variant lacking the GAS2 domain (Shot^ΔCTail^::GFP), or Shot Cterm::YFP, a C-terminal Shot fragment containing only the GAS2 domain (Fig. 3 A). Of the above constructs, we noticed that Shot^ΔCH1^::GFP, but not the others, caused pruning defects when overexpressed under two copies of the c4da neuron driver *ppk-GAL4* (Fig. S2).

**Figure 3.**
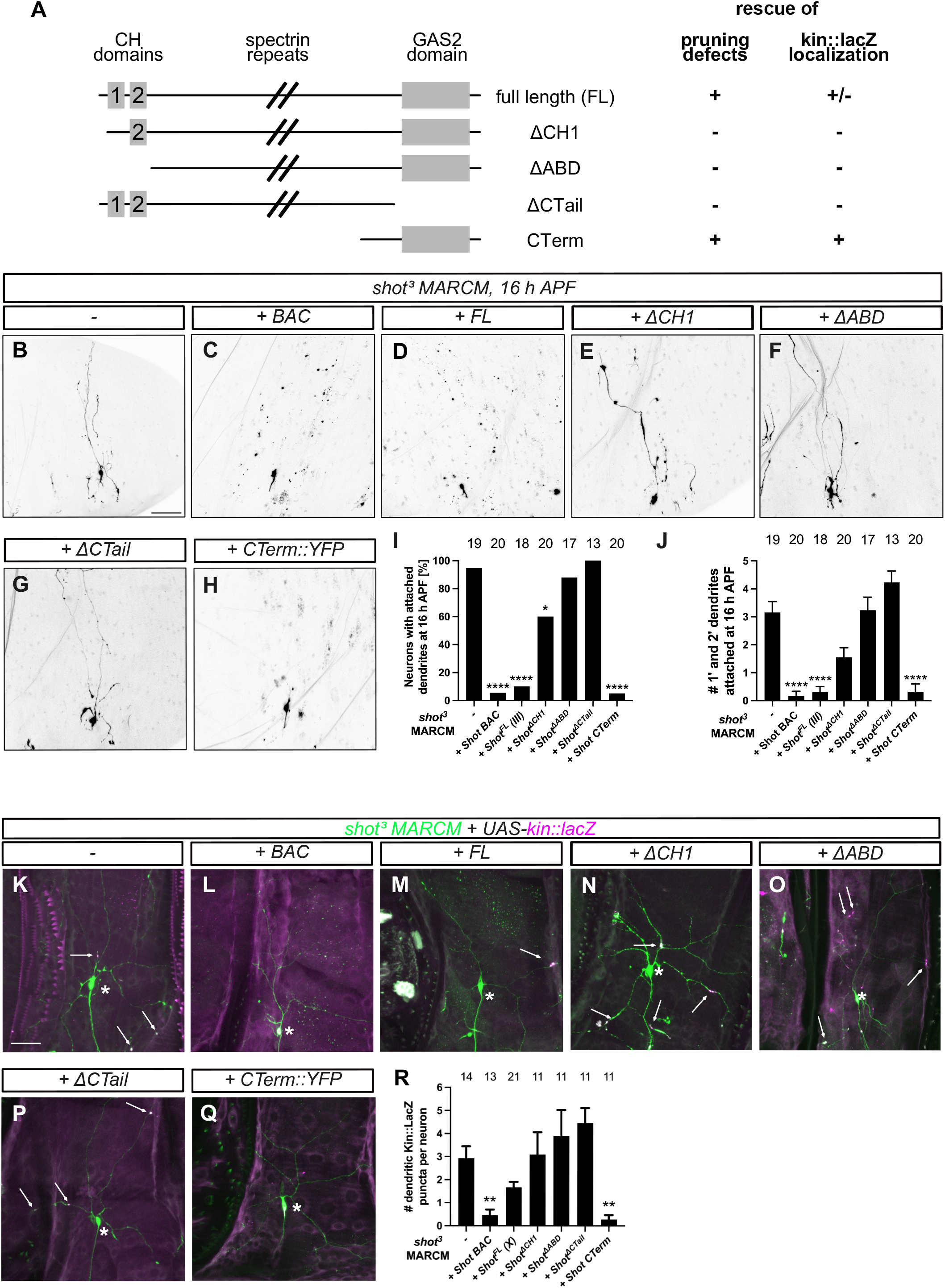
Shot domains important for pruning and microtubule organization. **A** Schematic of Shot domain structure and used UAS-Shot constructs. Right, summary of the phenotypic characterization below. CH, Calponin homology. **B** - **J** Ability of the indicated Shot UAS constructs to rescue the pruning defects of *shot^3^* mutant c4da neuron MARCM clones at 16 h APF. **B** *shot^3^* mutant c4da neuron. **C** *shot^3^*mutant c4da neuron rescued by a genomic Shot::YFP BAC construct. **D** *shot^3^* mutant c4da neuron expressing full-length Shot::GFP. **E** *shot^3^*mutant c4da neuron expressing Shot^ΔCH1^::GFP. **F** *shot^3^*mutant c4da neuron expressing Shot^ΔABD^::GFP. **G** *shot^3^* mutant c4da neuron expressing Shot^ΔCTail^::GFP. **H** *shot^3^* mutant c4da neuron expressing Shot C-term::YFP. **I** Penetrance of pruning defects. Number of neurons per sample is indicated above the graph. * P<0.05, **** P<0.0001, two-tailed Fisher’s exact test. **J** Number of primary and secondary dendrites attached to the cell body at the indicated timepoints. Values are mean +/- s. e. m., **** p<0.0001, Mann-Whitney U test. **K** – **R** Kin::lacZ localization in third instar *shot^3^* c4da neuron MARCM clones. **K** *shot^3^* mutant c4da neuron. **L** *shot^3^* mutant c4da neuron rescued by a genomic Shot::YFP BAC construct. **M** *shot^3^* mutant c4da neuron expressing full-length Shot::GFP. **N** *shot^3^*mutant c4da neuron expressing Shot^ΔCH1^::GFP. **O** *shot^3^*mutant c4da neuron expressing Shot^ΔABD^::GFP. **P** *shot^3^* mutant c4da neuron expressing Shot^ΔCtail^::GFP. **Q** *shot^3^* mutant c4da neuron expressing Shot C-term::YFP. Asterisks in **K** – **Q** denote soma position, and arrows mark dendritic kin::lacZ puncta. **R** Number of dendritic kin::lacZ puncta in third instar c4da neurons expressing the indicated Shot UAS constructs. Number of neurons per sample is indicated above the graph. Values are mean +/- s. e. m., ** p<0.01, Mann-Whitney U test. Scale bars in **B** and **K** are 50 μm.

Both the genomic Shot BAC construct and UAS-Shot^FL^::GFP rescued the pruning defects of *shot^3^* mutant neurons (Fig. 3 B - D, I, J). We next tested the domain mutants and found that while Shot^ΔCH1^::GFP slightly reduced the pruning defects of the mutant (Fig. 3 E, I, J), Shot^ΔABD^::GFP and Shot^ΔCTail^::GFP could not rescue them at all (Fig. 3 F, G, I, J). Strikingly, expression of Shot Cterm::YFP containing only the GAS2 domain led to a full rescue of the pruning defects (Fig. 3 H - J).

We next asked whether these domains are also important for Shot’s function in dendritic microtubule orientation. As the GFP tags of the Shot transgenes and the tdtomato fluorophore used to label MARCM clones precluded the use of fluorescently tagged EB1 constructs, we used kinesin-lacZ (kin::lacZ), a fusion between the kinesin-1 motor domain and β-galactosidase. Because the kinesin motor domain migrates towards microtubule plus ends, this fusion protein will always moves towards the soma in dendrites with normal plus end-in orientation and will thus show staining in the soma, but not dendrites. Consistent with the observation that loss of Shot causes microtubule orientation defects, kin::lacZ was depleted from the soma and found in dendritic puncta in *shot^3^* mutant c4da neurons (Fig. 3 K, R), but in animals carrying the genomic Shot BAC, it was localized in the soma (Fig. 3 L, R). UAS-based expression of Shot^FL^::GFP in mutant neurons did not fully rescue kin::lacZ localization but reduced the number of dendritic kin::lacZ puncta (Fig. 3 M, R). *shot^3^*mutant c4da neurons expressing Shot^ΔCH1^::GFP, Shot^ΔABD^::GFP or Shot^ΔCTail^::GFP did not confer rescue effects and showed many dendritic kin::lacZ puncta comparable to *shot^3^* mutant neurons (Fig. 3 N - P, R). Again, expression of Shot Cterm::YFP restored dendritic exclusion of kin::lacZ, and it could be seen in the soma (Fig. 3 Q, R).

Thus, in the context of full length Shot, both its actin and microtubule binding activities are required for its roles in dendrite pruning and microtubule orientation, but the isolated GAS2 domain alone was sufficient to rescue Shot function. This is reminiscent of a previous study reporting that full length Shot is in an autoinhibited state that can be relieved by actin binding (Applewhite *et al*, 2013). In support of the notion that microtubule orientation defects cause pruning defects, As the pruning and microtubule orientation defects of *shot* mutant neurons go hand in hand in our domain analysis.

### Shot cooperates with actin to promote microtubule orientation

To independently probe for a role for actin in dendritic microtubule organization, we sought to manipulate actin dynamics *in vivo*. As treatment with actin-depolymerizing drugs like latrunculin would likely be highly toxic to larvae, we considered Mical, a potent actin severing enzyme that breaks actin filaments by oxidizing actin on specific methionine residues (Grintsevich *et al*, 2017). Mical is highly specific for actin (Rajan *et al*, 2023) and its overexpression can disrupt actin bundles (Hung *et al*, 2010). Mical overexpression in larval c4da neurons did not disrupt dendrite morphology, but altered dendritic F-actin distribution at the first instar larval stage, as assessed by lifeact::GFP (Fig. S3). We next tested whether Mical overexpression has an effect on dendritic microtubule orientation and found that it caused a significantly increased number of EB1::GFP comets to move anterogradely (Fig. 4 A, B). When combined with Shot knockdown, Mical overexpression did not lead to an additive increase in the percentage of anterograde comets, potentially indicating epistasis between the manipulations (Fig. 4 A, B). However, we noted that the anterograde comets in the combined Shot knockdown/Mical overexpression neurons moved significantly faster than the retrograde comets, and also faster than the anterograde comets in the single manipulations (Fig. 4 C). Movement of the anterograde comets in these neurons was also more irregular than in other samples, as these comets frequently slowed down and then sped up again (Fig. 4 A).

**Figure 4.**
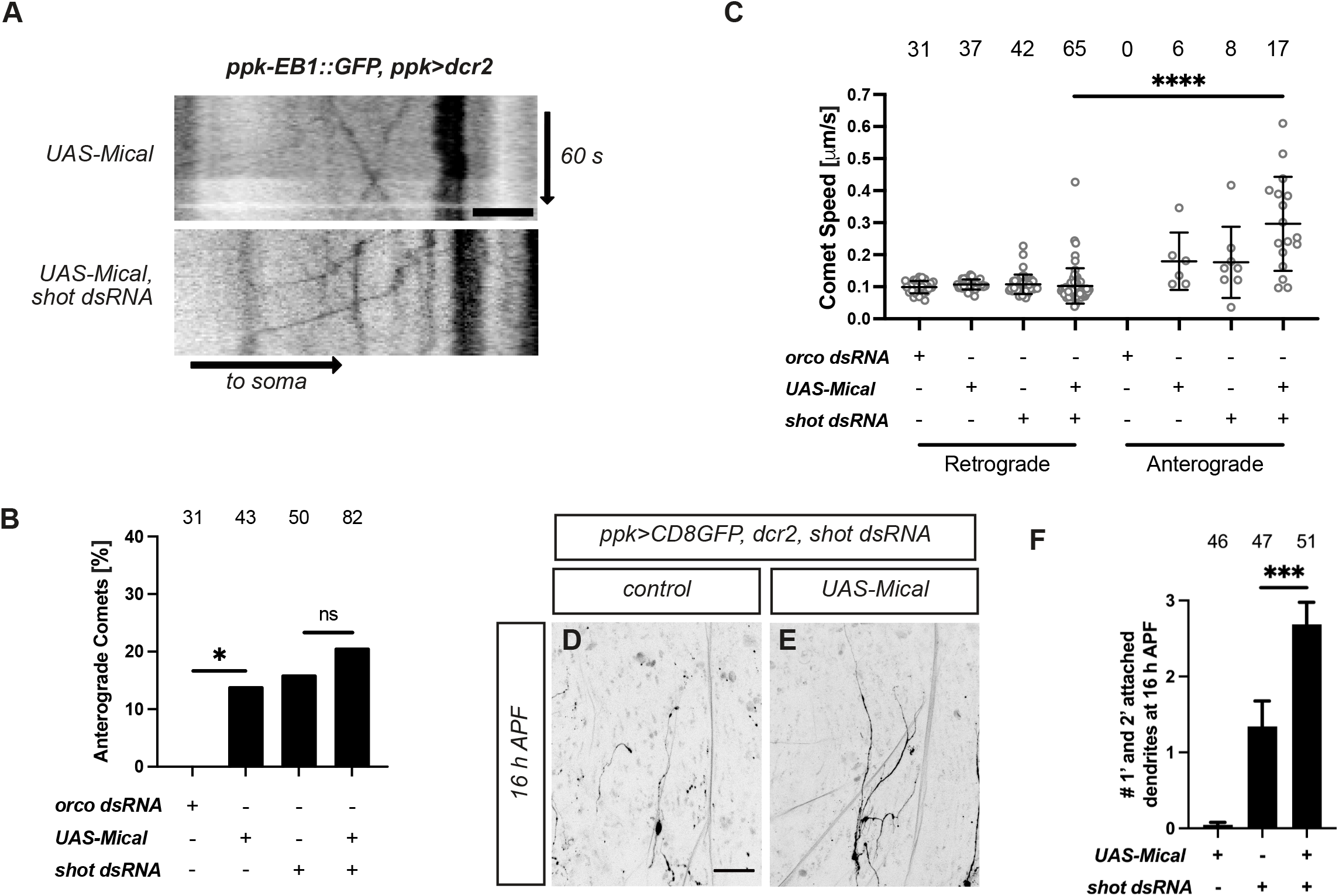
Shot and actin cooperatively regulate the behavior of dendritic microtubules. **A** Effect of actin severing on EB1::GFP comet movement. Mical was overexpressed in c4da neurons with or without Shot knockdown, and EB1 comets were analyzed as in Fig. 2 B. Upper panel, kymograph of EB1 comet movement in c4da neuron overexpressing Mical. Lower panel, kymograph of EB1 comet movement upon Mical overexpression and *shot* knockdown. **B** Penetrance of anterograde dendritic EB1 comets in control c4da neuron expressing *orco* dsRNA, or in neurons expressing *shot* dsRNA, Mical, or both (kymographs for the latter shown in **A**). Number of comets per sample is given in the graph. * P<0.05, Fisher’s exact test. **C** Speed of dendritic EB1 comets in **A**, **B**. **** P<0.0001, Wilcoxon’s test. **D** - **F** Synergistic effects of Shot knockdown and Mical overexpression on c4da neuron dendrite pruning. **D** C4da neuron expressing shot dsRNA and a control UAS construct at 16 h APF. **E** C4da neuron expressing shot dsRNA and UAS-Mical at 16 h APF. **F** Number of primary and secondary dendrites attached to the soma in c4da neurons overexpressing Mical, upon Shot knockdown, or both at 16 h APF. Number of neurons per sample is given in the graph. *** P<0.001, Mann-Whitney U test. Scale bars are 5 μm in **A** and 50 μm in **B**.

Conditions that cause mixed dendritic microtubule orientation also either cause c4da neuron dendrite pruning defects or enhance them (Herzmann et al., 2018). Mical overexpression alone does not cause pruning defects - in fact, it can suppress the pruning defects of certain ecdysone signaling mutants (Kirilly *et al*, 2009; Rode *et al*, 2018). However, we observed a synergistic enhancement of the pruning defects caused by Shot knockdown when we overexpressed Mical (Fig. 4 D - F). Thus, our data suggest that Shot and actin cooperatively regulate dendritic microtubule orientation.

### Shot is recruited to dendrite tips during early dendrite growth

As Shot is often recruited to specific subcellular locations via its actin binding domain (Nashchekin *et al*, 2024), we next assessed Shot localization in dendrites using GFP-tagged Shot constructs. We first visualized endogenously GFP-tagged Shot in third instar c4da neurons using structured illumination microscopy (SIM). In dendrite shafts, endogenously GFP-tagged Shot localized in closely spaced puncta close to the plasma membrane (Fig. 5 A), and transgenically expressed full length Shot (Shot^FL^::GFP) localized in a similar pattern (Fig. 5 B). We then visualized the two Shot variants lacking high affinity actin binding, Shot^ΔCH1^::GFP and Shot^ΔABD^::GFP, and found that they also localized close to the plasma membrane in a very similar pattern to full-length Shot (Fig. 5 C, D). Thus, actin binding is unlikely to be required for Shot cortical localization in dendritic shafts.

**Figure 5.**
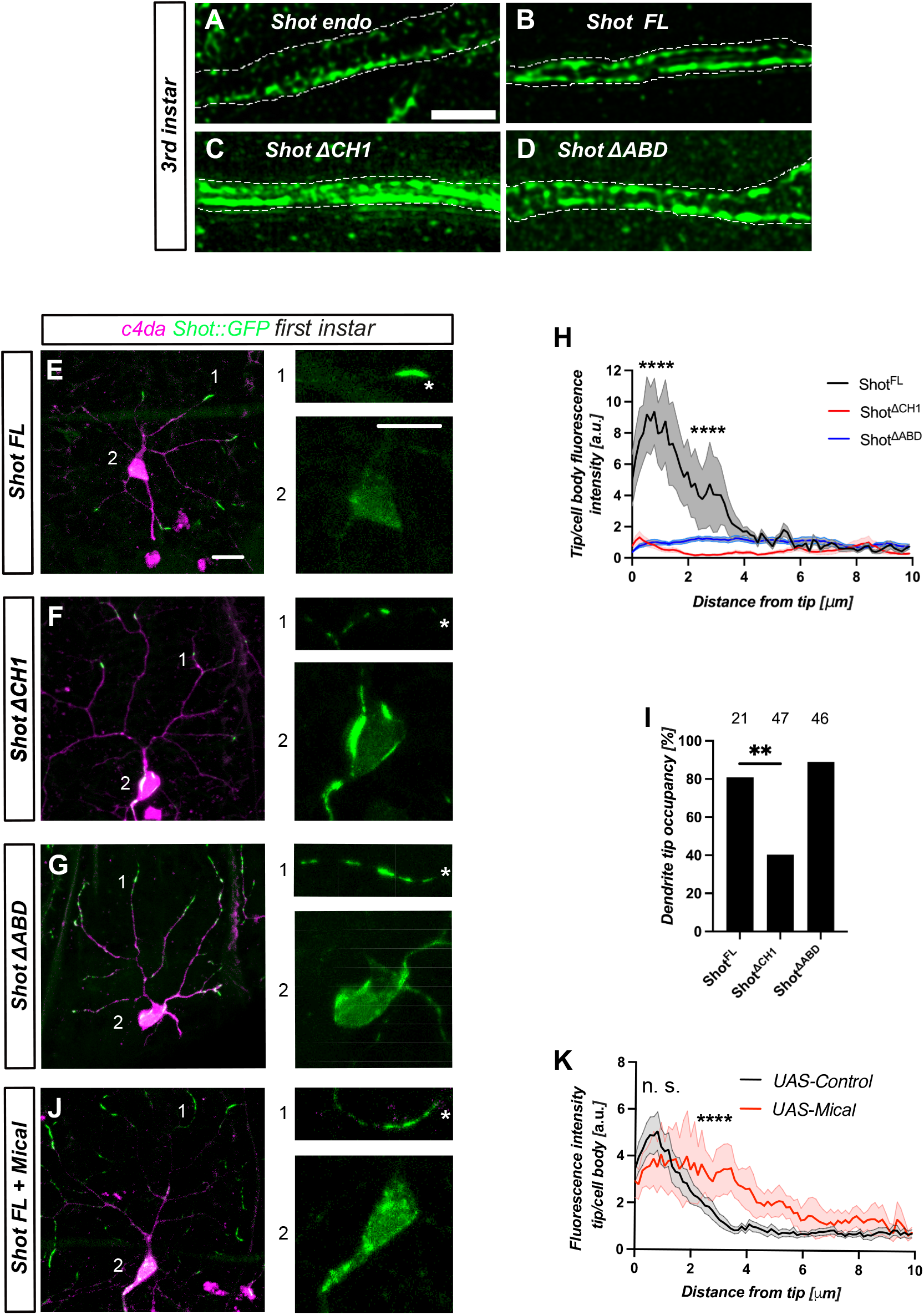
Shot is recruited to tips of growing dendrites via its actin binding domain. The indicated GFP-tagged Shot constructs and tdtomato were expressed in c4da neurons under *ppk-GAL4* and visualized at the indicated developmental stages. **A** - **D** Actin binding is not required for Shot localization in shafts of mature dendrites. The indicated Shot::GFP constructs were visualized in third instar primary c4da neuron dendrites by immunofluorescence followed by structured illumination microscopy. Borders of dendrites are indicated by white broken lines. **A** Endogenously tagged Shot::GFP. **B** Localization of transgenic full-length Shot (Shot^FL^::GFP) expressed under *ppk-GAL4*. **C** Localization of Shot^ΔCH1^::GFP expressed under *ppk-GAL4*. **D** Localization of Shot^ΔABD^::GFP expressed under *ppk-GAL4*. **E** - **K** Localization of the indicated Shot::GFP variants in first instar c4da neurons. Images on the right show close-ups of dendrite tip (1) and soma regions (2). Asterisks in the dendrite close-ups mark the position of the tip. **E** Localization of Shot^FL^::GFP. **F** Localization of Shot^ΔCH1^::GFP. **G** Localization of Shot^ΔABD^::GFP. **H** Fluorescence intensity profiles of Shot::GFP variants in the distal 10 μm of first instar dendrites in **E** - **G**. Intensity values are normalized to Shot::GFP intensity in the cell body. Solid lines indicate average, envelopes indicate s. d. (N=10 neurons). Compared to full length Shot, GFP intensities of Shot^ΔCH1^ and Shot^ΔABD^ in the 0-2 μm and 2-4 μm brackets from the tip were significantly reduced (P<0.0001, Mann Whitney U test). **I** Dendrite tip occupancy of Shot::GFP variants in **E** - **G**. ** P<0.01, Fisher’s exact test. **J**, **K** Effect of Mical overexpression on Shot^FL^::GFP localization. **J** Localization of Shot^FL^::GFP in a c4da neuron overexpressing Mical. **K** Fluorescence intensity profiles of Shot^FL^::GFP in the distal 10 μm of first instar dendrites with or without Mical overexpression. Shot::GFP intensity upon Mical overexpression was not significantly different from Shot::GFP without Mical in the 0-2 μm bracket, but highly significant in the 2-4 μm bracket (N=10 neurons, P<0.0001, Mann Whitney U test). Scale bars are 5 μm in **A** and 10 μm in **E** (larger image and close-up).

The control of dendritic microtubule organization can likely be divided in early establishment and later maintenance processes. To assess potential Shot functions at other developmental stages, we investigated the localization of transgenic Shot::GFP from the first larval instar until the early pupal stage (Fig. S4). Due to strong background from neighboring tissues, we were not able to visualize endogenously GFP-tagged Shot at all stages and throughout the whole c4da neuron. Strikingly, Shot::GFP was highly enriched at dendrite tips in first instar c4da neurons, but could only be detected at low levels in the soma and proximal dendrites (Fig. 5 E, H, I). From the second instar onwards, Shot::GFP appeared relatively evenly distributed in the soma and along both axons and dendrites (Fig. S4), indicating that Shot dynamically changes its localization during development. To test whether Shot dendrite tip localization depends on actin, we next imaged the actin binding-deficient variants Shot^ΔCH1^::GFP and Shot^ΔABD^::GFP in first instar c4da neurons. Shot^ΔCH1^::GFP accumulated in several dots per dendrite that could be many micrometers away from the tips (Fig. 5 F, H). Overall, significantly fewer distal dendrites were occupied by Shot^ΔCH1^::GFP compared to full length Shot (Fig. 5 F, I). Shot^ΔABD^::GFP was found at tips (Fig. 5 G, I), but always also localized in dots along dendritic shafts, such that there was no effective concentration at tips (Fig. 5 G, H). To independently test whether Shot tip localization is actin-dependent, we used UAS-Mical to disrupt actin. In c4da neurons overexpressing Mical, the Shot signal at tips was significantly broadened and Shot puncta could be seen along dendrite shafts (Fig. 5 J, K), similar to what was observed with Shot^ΔABD^::GFP. We conclude that actin binding is required to transiently recruit and concentrate Shot at dendrite tips during the early stages of c4da neuron dendrite growth.

### Shot acts early and is linked to end binding proteins

The observation that the Shot actin binding domain is required for both early dendrite tip localization and uniform dendritic microtubule orientation suggests that Shot function during early dendrite development is crucial for microtubule polarity. We therefore tested whether loss of Shot already causes defects in microtubule orientation during early development. When we imaged the directionality of EB1::GFP comets in first instar c4da neuron dendrites, we indeed found a significant increase in plus end-out microtubules upon loss of Shot (Fig. 6 A).

**Figure 6.**
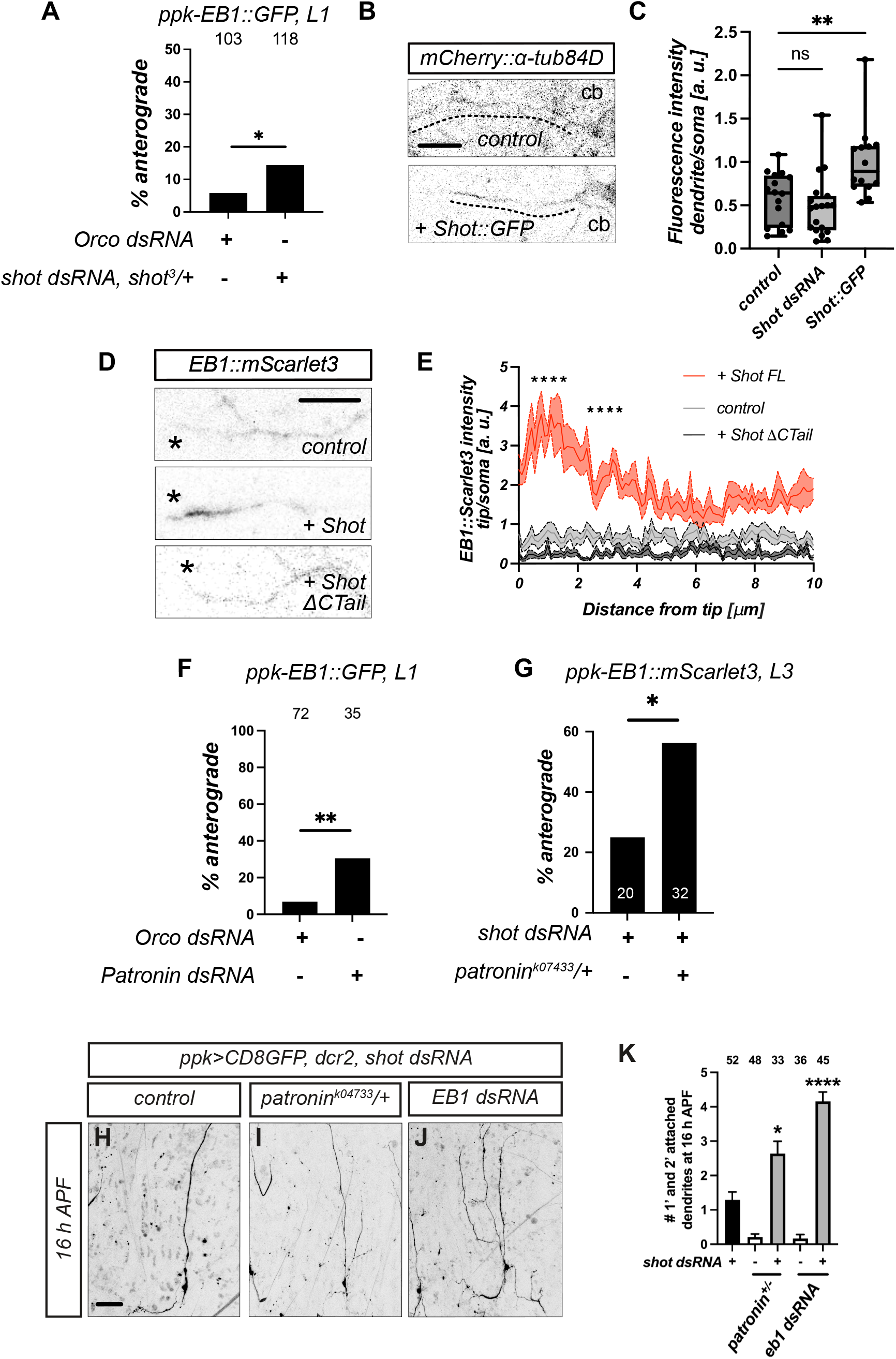
Shot acts during early development and interacts with end binding proteins. **A** Frequency of anterograde EB1 comets at the L1 stage in control (Orco dsRNA) and shot knockdown neurons (shot dsRNA, *shot^3^*/+). Number of comets per sample is shown in the graph, * P<0.05, Mann Whitney U test. **B** Microtubules in dendrites of first instar control neurons or neurons expressing Shot::GFP were visualized using mCherry-tagged α-tubulin in single confocal slices. **C** Dendritic microtubule density for samples in **B**. Dendritic mCherry-α-tubulin 84B fluorescence intensity was normalized to that of the soma. N= 14 - 22 dendrites from 9-14 neurons, * P<0.05, Student’s t-test. **D** Shot can recruit EB1 to dendrite tips. EB1::mScarlet3 was expressed in c4da neurons alone, with Shot^FL^::GFP, or with Shot^ΔCTail^::GFP, and visualized at the first instar larval stage. Positions of dendrite tips are indicated by asterisks. **E** Fluorescence intensity profiles of EB1::mScarlet3 in the distal 10 μm of first instar dendrites. Solid lines indicate average, envelopes indicate s. e. m. (N=9-10 dendrites, 3 neurons each). Significance between genotypes was calculated for 0 - 2 and 2 - 4 μm distances from the tip. **** P<0.0001, Mann Whitney U test. **F** Frequency of anterograde EB1 comets at the L1 stage in control (Orco dsRNA) and *patronin* knockdown neurons (patronin dsRNA). Number of comets is shown in the graph. ** P<0.01, Mann Whitney U test. **G** Frequency of anterograde EB1 comets at the L3 stage in *shot* knockdown neurons (*shot* dsRNA) and *shot* knockdown neurons in a heterozygous *patronin* mutant background. Number of comets is shown in the graph. * P<0.05, Mann Whitney U test. **H** - **K** Synergistic genetic interactions between *shot* and end binding proteins during c4da neuron dendrite pruning. **H** - **J** *shot* dsRNA was expressed in c4da neurons of animals in the indicated backgrounds, and pruning defects were quantified at 16 h APF. **H** C4da neuron expressing *shot* dsRNA. **I** C4da neuron expressing *shot* dsRNA in a *patronin*/+ background. **J** C4da neuron co-expressing *shot* and *EB1* dsRNAs. **I** Severity of pruning defects in **H** - **J** and in respective controls. N = 33 - 52 neurons, ** P<0.01, **** P<0.0001, Wilcoxon’s test. Scale bars are 5 μm in **B**, 2 μm in **D** and 50 μm in **E**.

To assess how Shot might affect microtubules at this stage, we next imaged mCherry-tagged α-tubulin live in first instar neurons and manipulated Shot levels. Interestingly, Shot overexpression caused a clear increase in microtubule density in first instar dendrites (Fig. 6 B, C). Shot knockdown did not cause a significant decrease of the dendritic microtubule signal in this experiment (Fig. 6 C), but this could in part be due to technical reasons, as we could not use the strongest Shot manipulation (Shot knockdown only, instead of knockdown in a heterozygous *shot* mutant background). To visualize dendritic microtubule structure at high resolution in third instar larval c4da neurons with the stronger manipulation, we also used stimulated emission depletion (STED) microscopy (Fig. S5). While microtubules appeared as several continuous and bundled filaments in controls, fewer filamentous structures were visible in dendrites of c4da neurons lacking Shot, the microtubule staining was more punctate, and the width of the microtubule bundles significantly thinner. Together with the domain analysis demonstrating the importance of the GAS2 domain, these data suggest that Shot promotes stabilization of dendritic microtubules. Shot is known to bind to the plus end binding protein EB1 via several SxIP motifs in its C-terminal region (Slep *et al*, 2005; Alves-Sila *et al*, 2012), and EB1 is also implicated in dendritic microtubule orientation (Mattie et al., 2010). In the presence of Shot::GFP, red fluorescently tagged EB1 also became recruited to dendrite tips at the first instar and colocalized with Shot::GFP (Fig. 6 D, E). This depended on the presence of the Shot C-terminal region including the SxIP motifs, as Shot^ΔCTail^ expression did not recruit EB1 (Fig. 6 D, E). Very rarely, EB1 comets could be seen emanating from a Shot/EB1-rich tip (Fig. S5).

The minus end binder Patronin also interacts with Shot (Nashchekin *et al*, 2016) and is required for uniform plus end-in dendritic microtubule orientation (Wang et al., 2019). We confirmed in immunoprecipitation experiments that a Shot fragment comprising spectrin repeats 27-32 interacts with Patronin (Fig. S5), similar to what has been reported for the mammalian Shot and Patronin homologues (Noordstra *et al*, 2016; Ning *et al*, 2016). Similarly to Shot, the mechanism of Patronin in dendritic microtubule orientation likely involves microtubule stabilization (by protecting minus ends from deolymerization) (Wang et al., 2019). Patronin knockdown caused an increase in the frequency of plus end-out microtubules already at the first instar stage (Fig. 6 F), and a heterozygous P element mutant in *Patronin* significantly increased the percentage of plus end-out microtubules at the third instar when combined with Shot knockdown (Fig. 6 G). We also tested for genetic interactions between Shot and end binding proteins using pruning as readout and found that both the heterozygous *Patronin* mutant and a dsRNA construct targeting EB1 strongly enhanced the pruning phenotypes seen upon shot knockdown (Fig. 6 H - K), again demonstrating the link between dendritic microtubule orientation and pruning. Thus, Shot likely cooperates closely with end binders in dendrites.

### Rab11 is linked to dendritic microtubule regulation in c4da neurons

A tip-based mechanism for microtubule orientation has first been proposed in *C. elegans* PVD sensory neurons, where Rab11-positive vesicles are part of a transient ncMTOC that localizes to the tips of growing dendrites (Liang *et al*, 2020; Harterink *et al*, 2018). We therefore asked whether a similar transient MTOC would also exist in Drosophila sensory neurons. In Drosophila sensory neurons, vesicle-based dendritic MTOCs have been proposed on either Golgi outposts (Ori-McKenney *et al*, 2012; Mukherjee *et al*, 2020; Yalgin *et al*, 2015) or Rab5 endosomes (Nguyen *et al*, 2014; Weiner *et al*, 2020), and microtubules have been observed originating in dendritic shafts, at branchpoints and at tips (Mukherjee *et al*, 2020; Yalgin *et al*, 2015; Weiner *et al*, 2020; Liang *et al*, 2020).

We and others have shown that Rab11 is required for c4da neuron dendrite pruning via an unclear mechanism (Krämer *et al*, 2019; Lin *et al*, 2020). To test whether Drosophila Rab11 interacts with microtubule regulators involved in pruning regulation, we used immunoprecipitation of tagged proteins from transfected S2 cells. This showed that Rab11 could be co-precipitated with Msps/XMAP215, a microtubule polymerase linked to pruning and dendritic microtubule organization (Tang *et al*, 2020) (Fig. 7 A). This interaction depended on GTP binding by Rab11, as GTP-binding-deficient Rab11 S25N showed a reduced interaction with Msps (Fig. 7 A). We next asked whether Rab11 colocalizes with potential MTOC components in c4da neuron dendrites using fluorescently tagged transgenes. At the first instar, Rab11::mCherry puncta could be seen overlapping with γ-tubulin::GFP both at dendrite branchpoints and tips (Fig. 7 B). To test whether such puncta could nucleate microtubules, we observed Rab11::mCherry together with EB1::GFP. In first instar neurons, a fraction of EB1::GFP comets could be observed originating at dendritic Rab11::mCherry puncta (6/9 neurons, 10/58 comets) in dendritic shafts or at branchpoints (Fig. 7 C, D). All of these comets moved retrogradely (10/10). Partially because of low EB1::GFP fluorescence, we could not detect dendrite tips - or comets originating from tips - in these experiments. In support of the idea that this Rab11/microtubule link might be transient, no EB1::GFP comets emanating from Rab11::mCherry puncta could be observed at the third instar stage (0/9 neurons, 27 comets) (Fig. 7 D). We next knocked down Rab11 and Rab5 and assessed the effects on microtubule orientation. Knockdown of Rab11, but not Rab5, caused a significant increase in the fraction of anterograde EB1 comets (Fig. 7 E, F) as well as increased comet speed at the third instar stage (Fig. 7 G). At the first instar stage, Rab11 knockdown caused a milder increase in anterograde comets that was not significant (Fig. S6). Thus, our evidence links Rab11 to dendritic microtubule regulation in Drosophila c4da neurons, but in part for technical reasons we cannot be certain whether it acts through the same tip-based mechanism as in *C. elegans* PVD.

**Figure 7.**
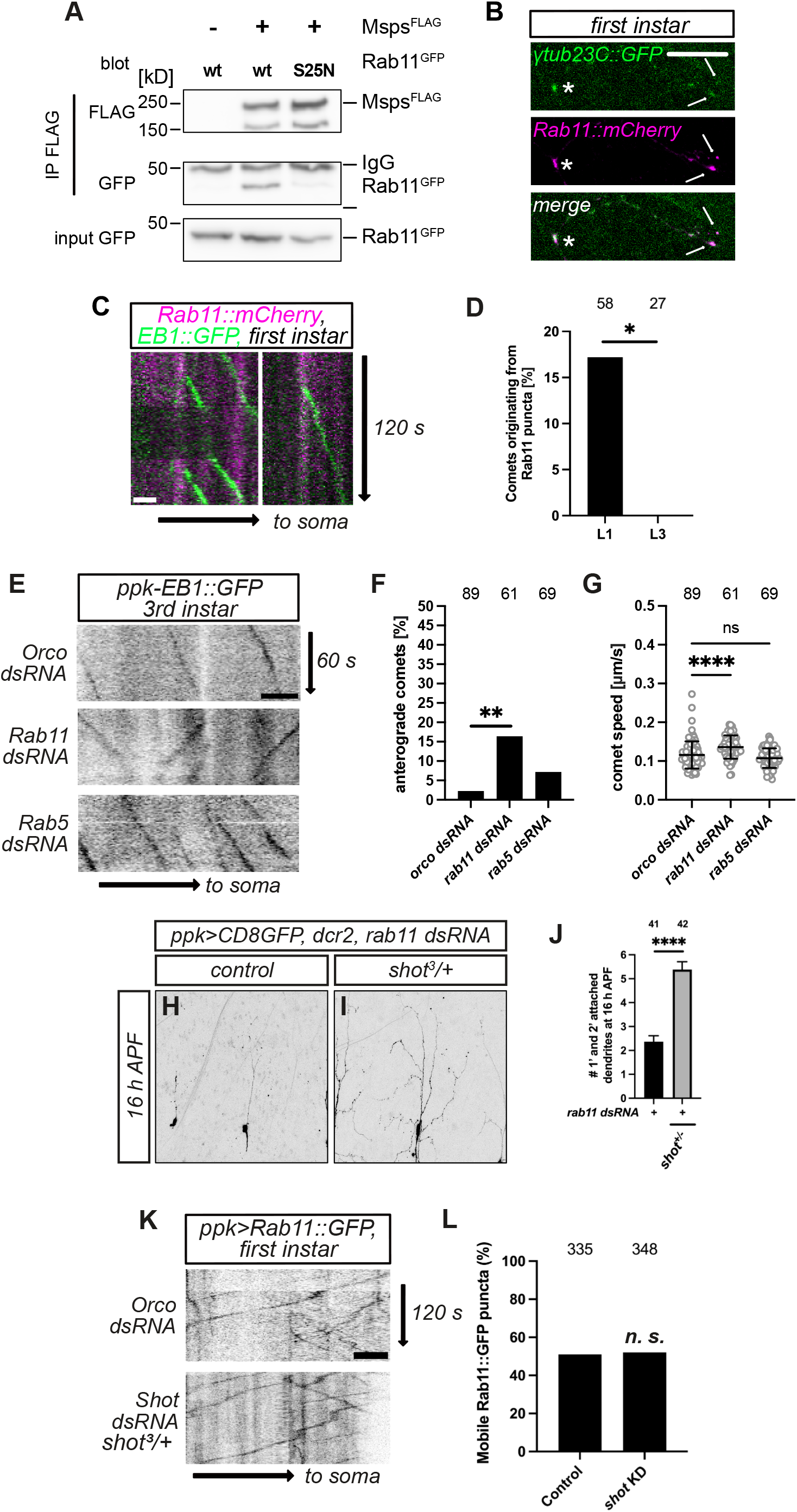
Evidence that Rab11 is linked to developmental dendritic microtubule regulation. **A** GFP-tagged Rab11 (wt or S25N) and FLAG-tagged Msps were cotransfected into S2 cells and immunoprecipitated with FLAG beads. Inputs and immunoprecipitates (IP) were blotted with the indicated antibodies. Sizes of molecular weight markers in kiloDalton (kD) are shown. IgG denotes antibody heavy chains. **B** Rab11 colocalizes with the MTOC marker γ-tubulin in growing c4da neuron dendrites. Rab11::mCherry and γ-tubulin23C::GFP were coexpressed under *ppk-GAL4* and visualized at the first instar. Arrows show Rab11/ γ-tubulin-positive puncta at dendrite tips, the asterisk denotes a double-positive dot at a branchpoint. **C** Microtubules can originate at Rab11 puncta in dendrites. Rab11::mCherry and EB1::GFP were coexpressed in c4da neurons, and EB1 comets were visualized at the first instar. Example kymographs show Rab11::mCherry (magenta) and EB1::GFP (green). **D** Percentage of EB1 comets arising from dendritic Rab11 puncta at the first and third instars. Number of comets is shown in the graph. * P<0.05, Fisher’s exact test. **E** Effect of Rab11 knockdown on dendritic microtubule orientation. Kymographs show EB1::GFP movement in third instar c4da neuron dendrites. Upper panel, control c4da neuron expressing Orco dsRNA; middle panel, c4da neuron expressing Rab11 dsRNA; lower panel, c4da neuron expressing Rab5 dsRNA. **F** Penetrance of anterograde comets in **D**. Number of comets in **E** and **F** is shown in the graph. ** P<0.01, two-tailed Fisher’s exact’s test. **G** Speed of EB1::GFP comets in **D**. **** P<0.0001, Wilcoxon’s test. **H** - **J** Rab11 and Shot interact genetically during dendrite pruning. *Rab11* dsRNA was expressed in c4da neurons of control animals (**H**) or in a *shot^3^*/+ heterozygous background (**I**), and neurons were visualized at 16 h APF. **J** Severity of pruning defects in **H**, **I**. N = 41 and 42 neurons. **** P<0.0001, Wilcoxon’s test. **K** Kymograph analysis of Rab11::GFP movement in dendrites of first instar control c4da neurons or *shot* knockdown neurons (lower panel). **L** Fraction of mobile Rab11::GFP particles in samples in **K**. Number of Rab11::GFP particles is shown in the graph. n. s., not significant, Fisher’s exact test. Scale bars are 5 μm in **B**, **C**, **E**, and **K**, and 20 μm in **H**.

As Shot has previously been shown to regulate Rab11 localization in epithelia (Khanal *et al*, 2016), we tested whether the two factors might also act in a common pathway in dendrites. We first used pruning phenotypic strength to assess a potential genetic interaction and found that Rab11 knockdown in a *shot^3^*/+ heterozygous background caused significantly more higher order dendrites to stay attached to the soma than with Rab11 knockdown alone, indicative of enhancement (Fig. 7 H - J). To observe potential effects in dendrites more directly, we expressed GFP-tagged Rab11 and visualized it using live microscopy at the first instar stage. Rab11::GFP vesicles in dendrites could be found along the whole dendritic shaft and were highly mobile (Fig. 7K). However, neither the mobility nor the overall localization of these puncta were changed upon shot knockdown (Fig. 7L). Thus, Shot is unlikely to directly regulate Rab11 trafficking in dendrites.

## Discussion

Microtubule orientation in neurites is crucial for neuronal polarity and developmental processes like neurite pruning, but the mechanisms that shape neuronal microtubule organization during development are still emerging. Here, we show that the spectraplakin Shot is required for the uniform plus end-in organization of dendritic microtubules. Our data suggest that Shot acts in a developmental timing-dependent fashion via localized microtubule stabilization at dendrite tips. Shot transiently localizes to dendrite tips in early growing dendrites. As this localization is actin-dependent, Shot recruitment may be mediated via a transient actin-rich growth cone-like structure. Importantly, the actin-binding domain required for tip recruitment is also required for correct microtubule orientation, indicating that Shot via a tip-based mechanism to orient microtubules. The main function of Shot at dendrite tips appears to be microtubule stabilization via the GAS2 domain. The surprising finding that this domain is sufficient to rescue Shot function in the absence of the large spectrin repeat region is consistent with the idea that Shot is regulated via autoinhibition, and actin binding may be required to disinhibit it (Applewhite *et al*, 2013). In addition to direct microtubule stabilization via the GAS2 domain, Shot can recruit the plus end binder EB1 (and potentially Patronin) to increase microtubule stability. This tip-based mechanism is reminiscent of the mechanism in the *C*. *elegans* PVD neuron, where a transient MTOC exists at tips of early growing dendrites. It was recently shown that in addition to bundling and stabilization, GAS2 domains can also nucleate microtubules *in vitro* (An *et al*, 2025). We think that it is unlikely that Shot itself acts to nucleate microtubules at dendrite tips because we were not able to detect a clear increase in microtubule nucleation from tips upon Shot overexpression. However, it is interesting to speculate that selective local microtubule stabilization at tips may already be sufficient to favour plus end-in polarity if the microtubules generated in other regions are sufficiently unstable, and microtubules at tips can only grow in the retrograde direction. This idea could be investigated in the future, e. g., through mathematical modelling. However, it is also possible that other, additional MTOC components localize to early dendrite tips in Drosophila sensory neurons as well. One candidate here is Rab11 which our analysis links to dendritic microtubule regulation, even though we did not see Rab11 enriched at tips. It is possible that the first instar stage, when most of our reagents start to work reliably, is already too late to visualize some of the very early growth events. In any case, our data suggest that early-stage tip-based mechanisms are a general feature of dendritic development.

Our data also do not rule out that Shot could anchor dendritic microtubules in mature third instar neurons, likely in cooperation with actin. Loss of Shot in combination with actin disassembly cause erratic comet behaviors (Fig. 4), which could be indicative of sliding, e. g., through defective microtubule anchoring (He *et al*, 2020). Alternatively, the observation that only the plus end-out oriented subset of comets is fast-moving could point to a different origin of these microtubules, e. g., they could enter the dendrites from the soma. Local membrane-bound microtubule stabilizers have been linked to the maintenance of dendritic microtubule orientation (He *et al*, 2020, Liang *et al*, 2025), and Shot might have a similar function. Crucially, both proposed functions of Shot in dendritic microtubule organization depend on actin binding, linking the two main cytoskeletal components in neurite organization.

Last but not least, our data again confirm our notion that uniform plus end-in dendritic microtubule orientation is required for c4da neuron dendrite pruning, as all Shot manipulations leading to a more mixed orientation caused pruning defects. As we do not have evidence for Shot regulation at the onset of the pupal stage (Fig. S4), the function of Shot crucial for pruning is most arguably its early role in setting up uniform microtubule orientation.

## Materials and Methods

### Fly Strains

C4da neurons were labeled by UAS-CD8::GFP or UAS-tdtomato expression under *ppk-GAL4* (Grueber *et al*, 2007). MARCM clones of *shot^3^*mutants (BL 5141) were induced with *SOP-FLP* (Matsubara *et al*, 2011) and labeled by tdtomato expression under *nsyb-GAL4^R57C10^*. Endogenously tagged Shot::GFP was from (Voelzmann *et al*, 2026) and the BAC carrying the SHOT genomic locus with a YFP tag was from (Nashchekin *et al*, 2016). UAS transgenes were UAS-Shot^RE^::GFP on the X chromosome (BL 29044), or mobilized to the third chromosome (this study), UAS-Shot^RC^::GFP (ΔCH1, BL 29042), UAS-Shot^ΔABD^::GFP (Hahn *et al*, 2021), UAS-Shot^ΔCTail^::GFP (Alves-Silva *et al*, 2012), UASp-Shot Cterm::YFP (Nashchekin *et al*, 2024), UAS-kin::lacZ (Clark *et al*, 1997), UAS-Mical (Terman *et al*, 2002), UAS-αtub84B::mCherry (Villars *et al*, 2022), UAS-Rab11::GFP (BL 8506), UAS-γtub23C::GFP (Nguyen *et al*, 2011), and UAS-cytoABKAR TA (Marzano *et al*, 2021) as inert UAS control, UAS-Brp^short^::Strawberry to label presynapses (gift from S. Sigrist). EB1 analyses were carried out using either UAS-EB1::GFP (Zheng *et al*, 2008), *ppk-EB1::GFP* (Arthur *et al*, 2015), UAS-EB1::mScarlet3 (gift from M. Rolls) or *ppk-EB1::mScarlet3* (this study). The following dsRNA lines were used: Orco (BL 31278) as control, Shot (BL 28336), Rab11 (VDRC 22198), Rab5 (VDRC 34096), EB1 (VDRC 24451). All dsRNAs were coexpressed with UAS-dcr2 (Dietzl *et al*, 2007). Additional mutant alleles were *patronin^k07433^* (Kyoto 111217).

### Cloning and transgenes

Shot sgRNAs GCTGCCCTCTCAGGCCGATT (target 1) and GAGTTCTCCAGAGTGGTCAC (target 2) were cloned into pCFD4w+ (gift from S. Schirmeier). For transgenesis, plasmids were injected into flies carrying the 86Fb acceptor site. For biochemical experiments, the Rab11 ORF was cloned into pUAST attB carrying a C-terminal GFP tag using TOPO/TA cloning, The S25N mutation was inserted by PCR mutatgenesis. N-terminally GFP-tagged Patronin was cloned into pUAST using standard restriciton sites. To express the Shot spectrin repeats 27-32, a fragment of the Shot ΟRF encoding amino acids 3626 - 4248 was cloned into pUAST carrying a C-terminal myc tag via TOPO/TA cloning. For Rab11::mCherry flies, the Rab11 entry clone was used to generate pUAST attB Rab11::mCherry, the insert was subcloned into pUAST, which was then injected into *w^1118^* flies using classical transposase-mediated transgenesis. To generate EB1::mScarlet3, a Drosophila codon optimized mScarlet3 (Gadella *et* al, 2023, synthesized by Twist Bioscience) was added after the EB1 coding sequence separated by a CACACCTCCACTACCGCTAGCAGGCCGGCCACGCGTGGTACCTTCTGGTCCA linker and cloned into pUAST. The resulting plasmid was injected into wild type embryos by Bestgene. The same insert was also inserted into the pDEST-APHIH vector carrying a *ppk* promotor fragment and injected into attP2 flies to generate *ppk-EB1::mScarlet3* flies. C-terminally FLAG-tagged Msps (isoform RB) was cloned into pUAST attB using standard cloning procedures.

### Immunofluorescence

Briefly, appropriately staged pupal filets were fixed in 4% formaldehyde, blocked in PBS with 0.3 % Triton X-100 and 10% goat serum, and incubated with antibody in blocking buffer over night. C4da neurons labeled by *ppk* promotor fusions, endogenous or transgenic GFP-tagged Shot were visualized with chicken (1:500, Aves labs), rabbit (1:1000, Invitrogen A11122) or mouse (1:1000, Invitrogen A1120) anti-GFP. Tdtomato and mcherry-tagged transgenes were detected with rat anti-mCherry (1:1000, Invitrogen) or rabbit anti-DsRed antibodies (1:1000, Clontech). Kin::β-galactosidase fusion proteins were detected with rabbit anti-β-galactosidase (Cappel, preabsorbed in-house, 1:700), and Drosophila futsch was detected with 22C10 (DSHB, 1:50). Secondary antibodies for regular immunofluorescence were conjugated to Alexa Fluor 488, 568 or 647. For STED microscopy of tagged tubulin, Abberior anti rabbit STAR RED were used as secondary antibodies.

### Microscopy, time lapse imaging

For analyses of dendrite pruning phenotypes at 16 h APF, animals were dissected out of the pupal case and dorsal ddaC c4da neurons in segments A2 - A5 were imaged live on a Zeiss LSM710 confocal microscope with a 20x Plan Apochromat water objective (1.0 NA). For regrowth analyses at 72 h APF, animals were dissected out of the pupal case and ventral v’ada c4da neurons in segments A2 - A4 were imaged as described (Sanal *et al*, 2023). For live protein localization analyses, first instar larvae were immobilized on double sided tape, third instar larvae were briefly anaesthesized with ether before imaging. Neurons were then imaged live on Zeiss LSM710 or LSM880 microscopes with 63x Plan Apochromat DIC M27 oil objectives (1.4 NA). For EB1::GFP or EB1::mScarlet3 live imaging, third instar larvae were anaesthesized with ether, and first instar larvae were immobilized on double-sided tape. Live analysis was performed on a Zeiss LSM880 microscope using a 40x Plan Apochromat FCS M27 (1.2 NA) oil objective. Consecutive images of a single plane were taken every second for 1 - 2 minutes, Z positions were adjusted if larvae moved. Movies were stabilized with the Fiji Image Stabilizer plugin, and the KymoResliceWide plugin was used to generate EB1 comet kymographs. Comet speed was read out manually from kymographs.

Immunofluorescence images of kin::lacZ were obtained on a Zeiss LSM 710 microscope using a 40x Zeiss C Apochromat water objective (1.1 NA). Structured illumination microscopy (SIM) was performed on a Zeiss Elyra 7 microscope with a 63x Plan Apochromat DIC M27 oil objective (1.4 NA) with an Edge 4.2 SIM camera (50 ms exposure time), and deconvolution was done using SIM^2^ software. All microscopes used ZEN Black software. Most Stimulated Emission Depletion Microscopy (STED) images were taken with an abberior FACILITY LINE (Abberior Instruments GmbH) equipped with an inverted IX83 microscope (Olympus), a 60x oil objective (Olympus UPLXAPO 60XO, NA 1.42), using pulsed excitation lasers at 640 nm and a pulsed STED laser operating at 775 nm and gated detection with avalanche photodiode element (APD) detectors, where acquisition operations were controlled by Lightbox Software (Abberior Instruments GmbH). Some 2D STED images were also taken with an abberior STEDYCON with an inverted IX83 microscope (Olympus), a 100x oil objective (Olympus UPLXAPO100XO, NA 1.45), using pulsed excitation lasers at 640 nm and a pulsed STED laser operating at 775 nm, continuous autofocus and gated detection with avalanche photodiode element detectors, where the acquisition operations were controlled by STEDYCON Software (abberior Instruments), and images were deconvolved using Huygens deconvolution software.

Microscopic images shown are maximum projections or (where indicated) single plane images. All processing and measurements were done in Fiji (Schindelin *et al*, 2012) using the plug-ins NeuronJ for dendrite length measurements.

### S2 cell culture, immunoprecipitation and Western blots

Appropriate pUAST plasmids (Rab11::GFP (wt or S25N), Msps^FLAG^, GFP::Patronin, myc::SRR27-32) were cotransfected into S2 cells with Actin5C-GAL4 as described (Herzmann *et al*, 2017). After 72 hours, cells were harvested in ice-cold PBS, and lysed in cold lysis buffer (100 mM NaCl, 5 mM MgCl_2_, 50 mM Tris/HCl pH 7.6, 5 % glycerol, 1 % Triton X-100, 1x complete protease inhibitor). Cleared lysates were incubated with anti-FLAG beads (Sigma A-2220) or with rabbit anti-GFP antibodies (Invitrogen A11222) bound to protein A sepharose (GE Healthcare 17-5280-01) for 2 hours. After three washes with lysis buffer, bound proteins were eluted in SDS sample buffer. Samples were run on 8 % gels and blotted with JL-8 anti-GFP (Clontech, 1:1000), FLAG M2 (Sigma F3165, 1:5000), or myc 9E10 (in house, 1:50) antibodies. Bands were visualized on an Amersham Imager 680.

### Quantification and statistical analyses

Phenotypic penetrance was assessed by counting the number of neurons with dendrites still attached to the soma. Here, significance was determined using a categorical two-tailed Fisher’s exact test. Length of unpruned dendrites were measured using the Fiji NeuronJ plugin and compared using the Wilcoxon Mann Whitney test. The number of dendritic kin::lacZ puncta was determined manually and compared using the Wilcoxon Mann Whitney test. EB1 comet orientation was assessed using a categorical two-tailed Fisher’s exact test, and a Mann Whitney test was used to compare comet speed. In order to quantify Shot or EB1 dendrite tip localization, intensity profiles of the distal 10 μm of each dendrite were averaged. The resulting curves were then sectioned into 2 μm segments, and data points from the individual intensity profiles for each segment were pooled and used for statistical comparisons between genotypes using the Mann Whitney test. To compare microtubule density, average cherry-tubulin fluorescence intensities along dendrites were normalized against cherry-tubulin fluorescence in the soma and compared using a Student’s t-test. Presynapse numbers were determined as described (Frommeyer *et al*, 2026). Two-sample t-tests were performed in Excel, all other tests were performed using Prism 10 software or online (Marx *et al*, 2016). Where necessary, data were corrected for multiple comparisons using Bennett’s test.

## Acknowledgments

We thank T. Zobel and S. Weischer at the Münster Imaging Network for expert advice on superresolution microscopy, D. Luchtman (abberior), S. Volkery and Nils Kirschnick (BioOptics facility, MPI for Molecular Biomedicine) for help with 2D and 3D STED, A. Prokop, G. Tavosanis, J. Wildonger, D. Nashchekin, S. Sigrist, the Bloomington, VDRC, and Kyoto stock centers and Addgene for fly lines and reagents. Requests for UAS-EB1::mScarlet3 should be directed to Melissa Rolls. We thank R. Hube for technical help in the inital stages of the project, T. Klein-Höing for help with cloning and A. Ziegler for advice on early larval c4da neuron imaging. MD, NS, SMF and NW are members of the CiM IMPRS and CRC1348 graduate schools, respectively. This work was supported by DFG grants RU1673/4-1, RU1673/6-1 and RU1673/10-1 to SR. The authors declare no competing financial interests.

## Author Contributions

M. Davies designed experiments, performed phenotypic analyses and contributed reagents. N. Sanal, N. Wolterhoff and S. M. Frommeyer performed phenotypic analyses. U. Gigengack performed biochemical experiments and contributed reagents. Y. Shen and I. Hahn contributed reagents. S. Rumpf designed experiments, performed biochemical analyses and contributed reagents. S. Rumpf wrote the manuscript with input from all other authors.

## Supplementary Figure legends

**Figure S1 (related to Fig. 1). Effect of microtubule regulators on c4da neuron synapse pruning and dendrite regrowth. A** - **C** Loss of Shot does not cause presynapse pruning defects. C4da neuron presynapses were visualized with *UAS-Brp^short^::Strawberry*, and presynaptic regions of control (Orco dsRNA) or Shot knockdown c4da neurons in the VNC (segments 3 - 5) were imaged at 24 h APF. **A** Control animal expressing Orco dsRNA in c4da neurons. **B** Animal expressing shot dsRNA in c4da neurons. **C** Number of Brp puncta in **A**, **B**. N=7-9 animals. Boxes show median, whiskers are s. d., not significant, Student’s t-test. **D** - **H** Loss of microtubule regulators causes dendrite regrowth defects. The indicated factors were knocked down in c4da neurons under *ppk-GAL4*, and neurons were imaged at 72 h APF. **D** Control c4da neuron expressing Orco dsRNA (N = 22 neurons). **E** C4da neuron expressing patronin dsRNA (N = 12 neurons). **F** C4da neuron expressing Shot dsRNA (N = 10 neurons). **G** C4da neuron expressing EB1 dsRNA (N = 10 neurons). **H** Total dendrite length in **D** - **G**. Values are mean +/- s. d., *** P<0.001, **** P<0.0001, Wilcoxon’s test. Scale bars are 20 μm in **A** and 50 μm in **D**.

**Figure S2 (related to Fig. 3). Effect of Shot domain mutant overexpression on c4da neuron dendrite pruning. A** - **E** Full length or truncated Shot variants were overexpressed in c4da neurons, and the effects on dendrite pruning were assessed at 16 h APF. **A** Control c4da neuron. **B** C4da neuron overexpressing full length Shot::GFP. **C** C4da neuron overexpressing Shot^ΔABD^::GFP. **D** C4da neuron overexpressing Shot^ΔCH1^::GFP. **E** C4da neuron overexpressing Shot Cterm::YFP. **F** C4da neuron overexpressing Shot^ΔCTail^::GFP. **G** Severity of pruning defects in **A** - **F**. Number of neurons per sample is shown in the graph, values are mean +/- s. e. m., **** P<0.0001, Wilcoxon’s test. The scale bar in **A** is 50 μm.

**Figure S3 (related to Fig. 4). Effect of Mical overexpression on dendritic actin distribution.** Actin was visualized at the first or third instar stages using LifeAct::GFP expressed under *ppk-*GAL4. **A** Control c4da neuron expressing Orco dsRNA at the first instar. **B** C4da neuron expressing UAS-Mical at the first instar. **C**, **D** LifeAct::GFP intensity in proximal (**C**) or distal (**D**) dendrites in **A**, **B** was normalized to soma intensity. Number of neurons is shown in the graph, values are mean +/- s. d., * P<0.05, Wilcoxon’s test. **E** Control c4da neuron expressing Orco dsRNA at the third instar. **F** C4da neuron expressing UAS-Mical at the third instar. **G**, **H** LifeAct::GFP intensity in proximal (**G**) or distal (**H**) dendrites in **E**, **F** was normalized to soma intensity. Number of neurons is shown in graph, values are mean +/- s. d., not significant, Wilcoxon’s test. Scale bars are 10 μm in **A** and 20 μm in **E**.

**Figure S4 (related to Fig. 5). Shot dendritic tip enrichment is transient.** Shot^FL^::GFP was expressed in c4da neurons under *ppk-GAL4* and c4da neurons were visualized live at the first, second and third instars, and by immunofluorescence at 5 h APF. **A** Shot::GFP localization in a first instar c4da neuron. Arrows denote the positions of dendrite tips. **B**, **B’** Shot::GFP localization in second instar c4da neurons. Panel **B** shows soma and proximal dendrites, **B’** shows distal dendrites and dendrite tips. **C**, **C’** Shot::GFP localization in third instar c4da neurons. **C**, soma and proximal dendrites, **C’**, distal dendrites and dendrite tips. **D** Shot::GFP localization in a c4da neuron at 5 h APF. All scale bars are 20 μm.

**Figure S5 (related to Fig. 6). Shot affects microtubule appearance and interacts with tip binding proteins. A** Microtubules in c4da neuron primary dendrites were visualized by immunofluorescence of mCherry::α-tubulin 84Β (under *ppk-*GAL4) followed by 2D STED. Upper panel, control c4da neuron expressing Orco dsRNA. Lower panel, shot knockdown in heterozygous *shot^3^*/+ animal. **B** Quantification of microtubule signal width in **A**. Intensity profiles of cross sections were taken, and width was measured at approximately 30 % maximum intensity. N=20-21 dendrites from 8-10 neurons, * P<0.05, Mann Whitney test. **C** Kymograph of EB1::mScarlet3 and Shot::GFP at a dendritic tip. Tip position is to the left, and arrows in the EB1::mScarlet3 and merge panels indicate a comet originating there (N=1). **D** A Shot fragment comprising spectrin repeats 27 - 32 interacts with Patronin. GFP::Patronin and myc-tagged Shot SRR27-32 were coexpressed in S2 cells and immunoprecipitated with antibodies against GFP. Inputs and immunoprecipitates (IP) were blotted with the indicated antibodies. Sizes of molecular weight markers in kiloDalton (kD) are shown. Scale bars are 2 µm in A and 5 µm in C.

**Figure S6 (related to Fig. 7). A** Effect of Rab11 knockdown on dendritic microtubule orientation at the first instar stage. Comet orientation was determined in first instar c4da neuron dendrites using *ppk-EB1::GFP* as in Figure 6. Shown is percentage of anterograde comets. Number of comets is shown in the graph, n. s., not significant, two-tailed Fisher’s exact’s test.

