## Supplementary figures and images for "Spectraplakin is required for dendritic microtubule organization via a tip-based mechanism"

### Supplementary Figs 1-6

#### Davies Figure S1

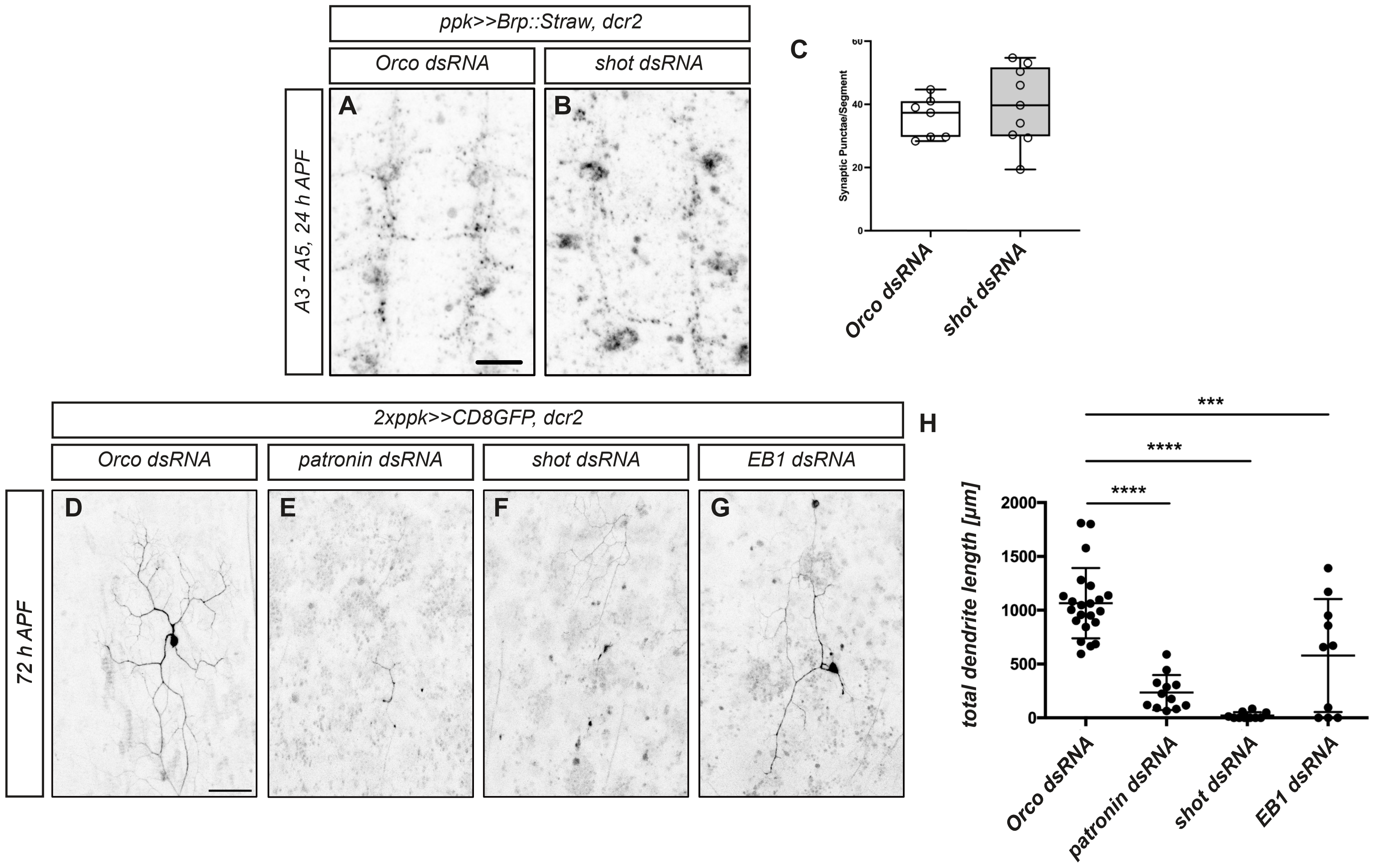

#### Davies Figure S2

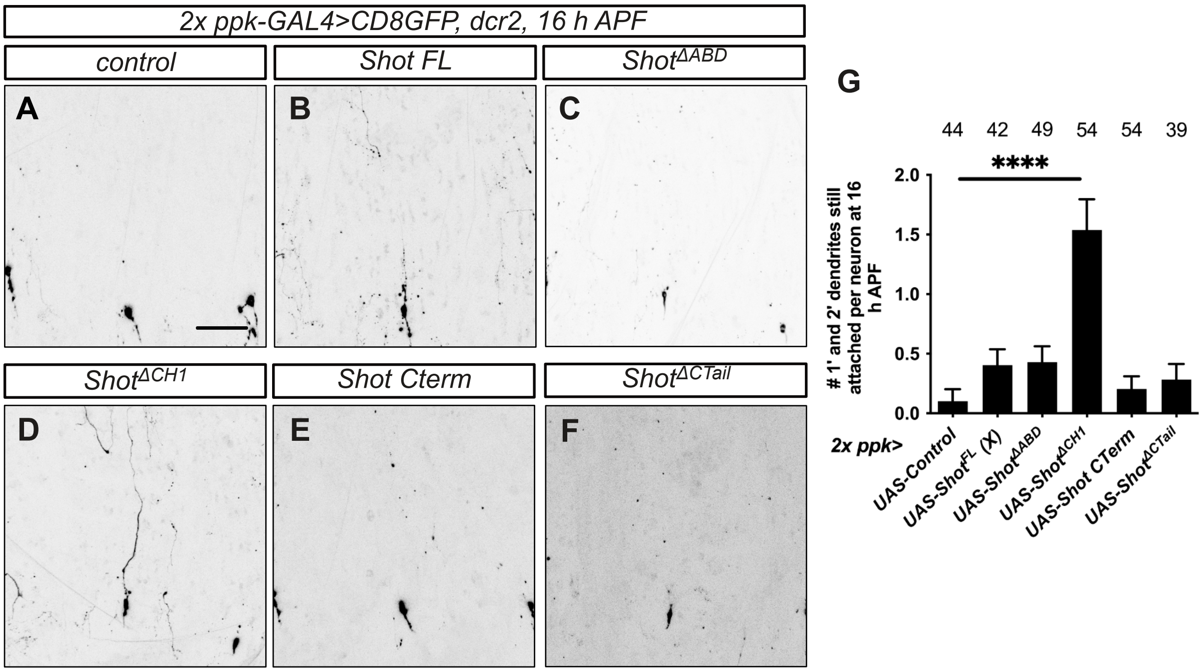

#### Davies Figure S3

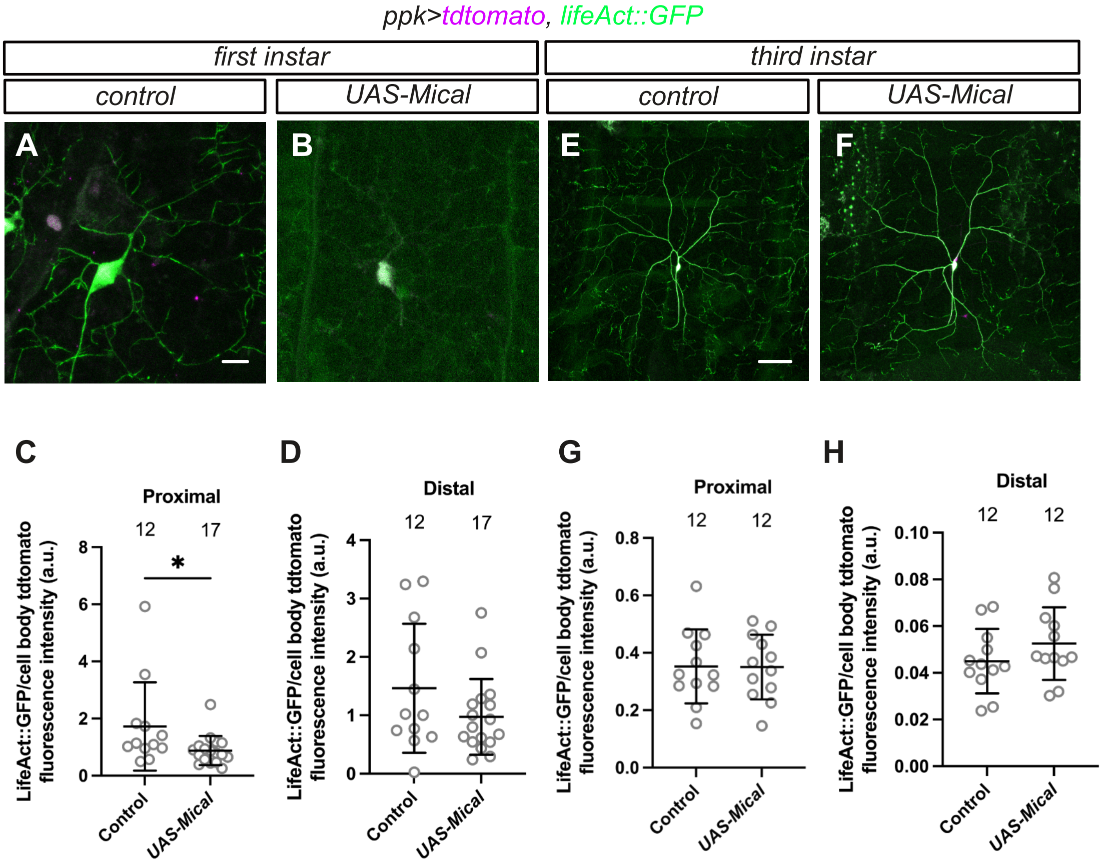

#### Davies Figure S4

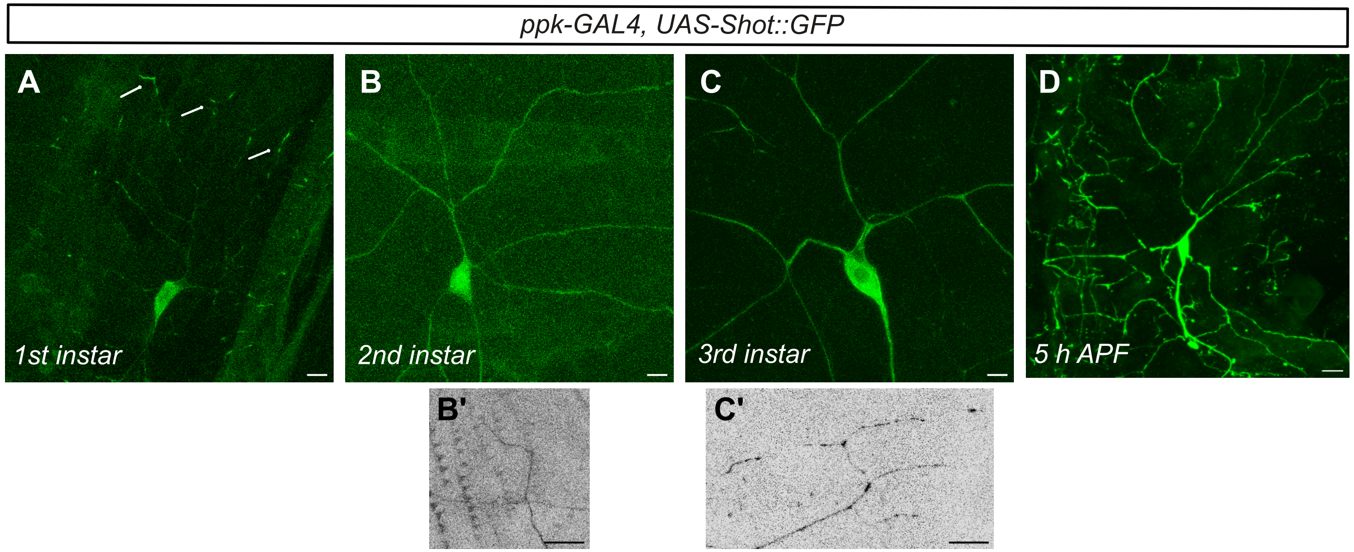

#### Davies Figure S5

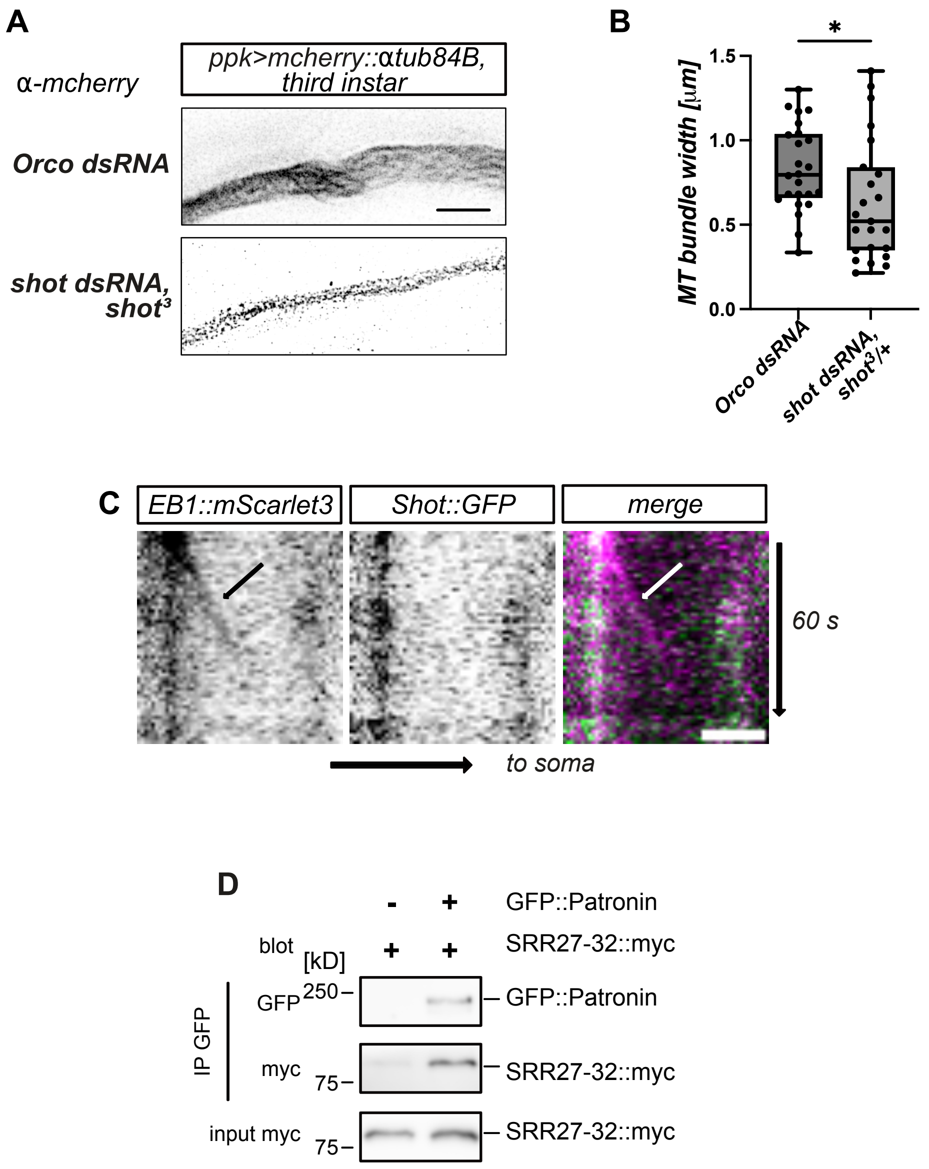

#### Davies Figure S6

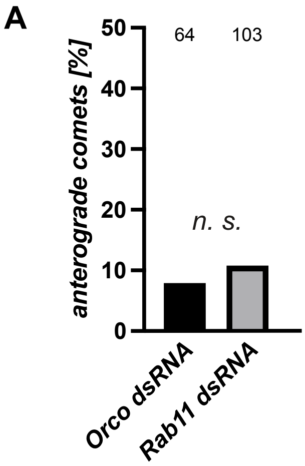
